# Elevator motion of chloride ion transporter SLC26A9 induced by STAS domain

**DOI:** 10.1101/2023.07.30.551137

**Authors:** Satoshi Omori, Yuya Hanazono, Hafumi Nishi, Kengo Kinoshita

**Affiliations:** Graduate School of Information Sciences, Tohoku University, Sendai, Miyagi 980-8579, Japan; Research Organization for Nano and Life Innovation, Waseda University, Shinjuku-ku, Tokyo 162-0041, Japan; Medical Research Institute, Tokyo Medical and Dental University, Bunkyo-ku, Tokyo 113-8510, Japan; Faculty of Core Research, Ochanomizu University, Tokyo 112-8610, Japan; Tohoku Medical Megabank Organization, Tohoku University, Sendai, Miyagi 980-8573, Japan; Institute of Development, Aging, and Cancer, Tohoku University, Sendai, Miyagi 980-8575, Japan

**Author notes:** These authors contributed equally to this work (Satoshi Omori, Yuya Hanazono).

## Abstract

The anion exchanger SLC26A9, consisting of the transmembrane (TM) domain and the cytoplasmic STAS domain, plays an essential role in regulating chloride transport across cell membranes. Recent studies have indicated that C-terminal helices block the entrance of the putative ion transport pathway. However, the precise functions of the STAS domain and C-terminal helix, as well as the underlying molecular mechanisms governing the transport process, remain poorly understood. In this study, we performed molecular dynamics simulations of three distinct models of human SLC26A9: full-length (FL), STAS domain removal (ΔSTAS), and C-terminus removal (ΔC), to investigate their conformational dynamics and ion binding properties. Stable binding of ions to the binding sites was exclusively observed in the ΔC model in these simulations. Comparing the FL and ΔC simulations, the ΔC model displayed enhanced motion of the STAS domain. Furthermore, comparing the ΔSTAS and ΔC simulations, the ΔSTAS simulation failed to exhibit stable ion bindings to the sites despite the absence of the C-terminus blocking the ion transmission pathway in both systems. These results suggest that the removal of the C-terminus not only unblocks the access of ions to the permeation pathway but also triggers STAS domain motion, gating the TM domain to promote ions’ entry into their binding site. Further analysis revealed that the asymmetric motion of STAS domain leads to the expansion of the ion permeation pathway within the TM domain, resulting in the stiffening of the flexible TM12 helix near the ion binding site. This structural change in the TM12 helix stabilizes chloride ion binding, which is essential for SLC26A9 elevator motion. Overall, our study provides new insights into the molecular mechanisms of SLC26A9 transport and may pave the way for the development of novel treatments for diseases associated with dysregulated ion transport.

**SIGNIFICANCE:** We explored the mechanism by which the human protein SLC26A9 transports chloride in the cell. SLC26A9 is a potential therapeutic target for patients with cystic fibrosis, as by targeting drugs to it, it may be possible to restore chloride ion transport in epithelial cells. To design therapeutic drugs, it is essential to understand how the protein works. Our findings support an elevator-type mechanism, in which chloride ions bind to SLC26A9 inside the cell and are then released by the protein to the extracellular environment. We find that the STAS domain of SLC26A9 has critical roles in binding chloride and induces conformational changes in the transmembrane domain that facilitate chloride transport.

## INTRODUCTION

Chloride ions, the most abundant anions in the extracellular fluid, are involved in a variety of physiological processes, such as the regulation of cellular pH, the control of membrane excitability of nerve or muscle cells, and epithelial transport (1-4). The dysfunction of chloride ion transport results in diverse disorders, including epilepsy, myotonic disorders, and cystic fibrosis (CF). CF is a genetic disorder caused by mutations in a chloride channel, cystic fibrosis transmembrane conductance regulator (CFTR) (5-7). Due to a low transport capacity for chloride ions, sticky secretions clog the bronchial and digestive tracts, leading to diseases such as pneumonia and bronchitis. Though CF is a serious disease frequently seen in Caucasians, there is no cure for CF at present, although there are therapeutics directed to CFTR with some efficacy, depending on the underlying mutation.

SLC26A9, which is a member of the Solute Carrier family 26 (SLC26) of anion transporter/channel proteins, is mainly expressed in the lung and gastric epithelium and contributes to mucociliary clearance and gastric acid production (8-11). SLC26A9 is regarded as a chloride ion channel or an uncoupled fast chloride transporter with channel-like activity (12-16). SLC26A9 interacts with CFTR to regulate chloride ion conductance (14,17). Variants of SLC26A9 are associated with CF-like lung disease (18,19) and modulate the response of the airways to CFTR-directed therapeutics (20,21). Given their colocalization and the functional correlation between SLC26A9 and CFTR, SLC26A9 is a potential therapeutic target for CF patients to restore chloride ion transport in epithelial cells.

SLC26 proteins belong to the Sulfate Permease (SulP) family that is conserved in various species from bacteria to mammal (22,23). The SulP family proteins consist of two domains: an alpha-helical transmembrane domain, and a STAS domain located in the cytoplasmic region. The topology of the transmembrane region consists of 14 transmembrane helices characterized by 7+7 inverted repeat folds (24-26). Recently, several high-resolution structures of SLC26A5 (27-29) and SLC26A9 (30,31) have been reported using cryo-electron microscopy (cryo-EM). The structures of mouse (30) and human (31) SLC26A9 were determined at 3.9 Å and 2.6 Å resolution, respectively. The cryo-EM structure of mouse SLC26A9 was performed using a truncated variant lacking the intrinsically disordered region of the long intervening loop in the STAS domain and C-terminus. The cytoplasmic STAS domain is important for forming a homodimeric structure and acts as a platform for interaction of subunits. Dimeric interactions exist not only between the transmembrane domains or STAS domains but also between the cytoplasmic surface of the transmembrane domain and the STAS domain of the opposite chain (Figure 1b). The determination of the structure of human SLC26A9 was performed using the full-length protein, although the long intervening loop in the STAS domain is disordered. The overall structure of human SLC26A9 is almost the same as that of mouse SLC26A9. Interestingly, the electron densities of the C-terminal regions are observed at the intracellular vestibule of the putative ion transport pathway and the C-terminal helices are bound near the entrance of the pathway (Figure 1a). The electrophysiological analysis showed that the C-terminal helices inhibit the transport of chloride ions. In addition, molecular dynamics (MD) simulations demonstrated that chloride ions approach the binding site through the pore region between the core and gate domains (31).

**FIGURE 1.**
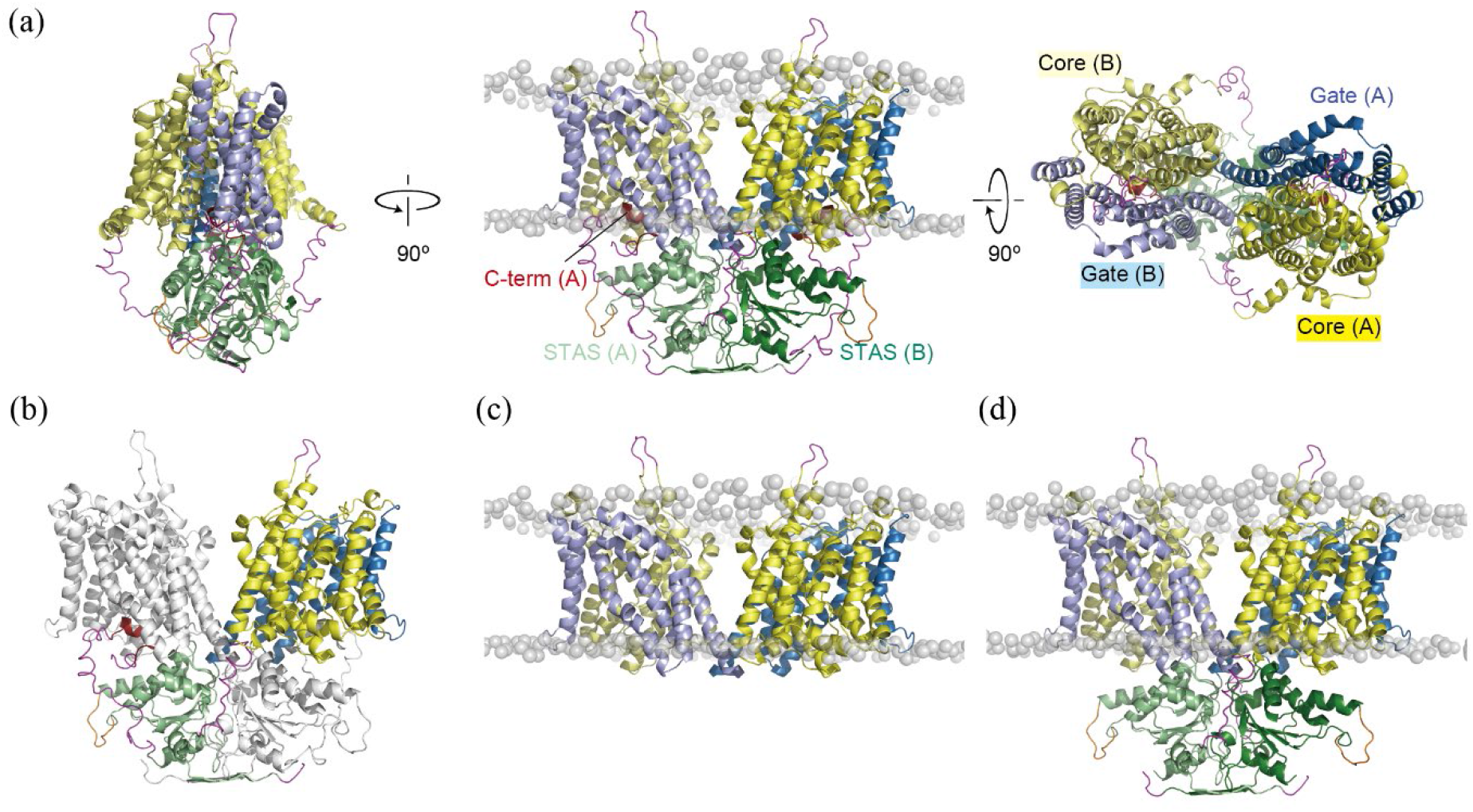
Initial structures of (a, b) FL, (c) ΔSTAS, and (d) ΔC simulations. Cartoon representations of core (yellow), gate (sky blue), STAS (light green) of chain A, and core (pale yellow), gate (light blue), STAS (deep green) of chain B are shown. C-terminal ordered regions (dark red), the modeled loops of the truncated portion (orange), and the short disordered regions (magenta) are also shown. Phosphorus atoms in the head groups of lipid membranes are shown as gray transparent spheres. (b) Chain B is colored white to show the characteristic X-shaped dimer structure.

Though the flexibility of the transmembrane helices that interact with the substrate has been implicated in transport movement (32), the series of movements before the elevator-like transport pathway, which can be a key feature of the transport mechanism of SLC26A9, is unclear. In addition, mutations in the STAS domain affect the transport function around the interface between the transmembrane regions (33-37), but little is known about the role of the STAS domain in chloride ion transport.

In order to better understand the dynamic properties of SLC26A9 and its role in chloride ion transport, we performed molecular dynamics (MD) simulations of full-length (FL), STAS domain removal (ΔSTAS), and C-terminus removal (ΔC) models of the human SLC26A9 protein. By analyzing the simulation data, we aimed to investigate the interactions between the STAS domain and the transmembrane region, as well as the effect of the C-terminal helices on chloride ion transport. Our simulations allowed us to observe the stable binding of chloride ions consistent with the elevator-like transport pathway. Moreover, our results revealed large and asymmetric motions of the STAS domain that promote the elevator motion by inducing conformational changes in the transmembrane domain. The insights gained from our study have important implications for the rational design of new drugs and more effective therapeutic treatments for disorders related to chloride ion transport, such as cystic fibrosis.

## MATERIALS AND METHODS

### Model building

To investigate the effect of the STAS domain and C-terminal helix on chloride ion permeation in SLC26A9, we constructed three models of the protein: full-length (FL), STAS domain removal (ΔSTAS), and C-terminus removal (ΔC) (Figure 1). We obtained the cryo-EM structure of human SLC26A9 from the Protein Data Bank (38) (PDB ID: 7CH1). However, since the intrinsically disordered region of the long intervening loop (residues 568-652) in the STAS domain is disordered in the cryo-EM structure of human SLC26A9, we modeled the truncated portion by replacing it with a loop consisting of residues 568-573 and 648-652 strung together. Additionally, we modeled the short disordered regions, which include residues 1-3, 27-44, 151-157, 742-772, and 785-791, to construct the FL model of residues 1- 791. We used CHARMM GUI (39) to model all disordered regions except residues 151-157, which we modeled using MODELLER ver. 9.25 (40). We built the ΔC model by excluding the C-terminal loop and residues 742-791, which include the loop region before and after the C-terminal loop. The ΔSTAS model was created from the ΔC model by excluding residues 1- 44 and 500-741 that correspond to the STAS domain.

### MD simulations

We inserted each SLC26A9 model (FL, ΔSTAS, and ΔC) into a 1-palmitoyl-2-oleoyl-sn-glycero-3-phosphocholine (POPC) model lipid membrane and immersed them in rectangular boxes filled with TIP3P (41) water molecules together with sodium and chloride ions, resulting in a salt concentration of 150 mM. The resulting systems contained 276,960, 209,084, and 277,656 atoms for the FL, ΔSTAS, and ΔC models, respectively. We built all systems using the Membrane builder (42) in CHARMM-GUI (39,43). We performed simulations using the MD program GROMACS ver 2018.2 (44). We employed the CHARMM36m force field (45) for proteins and lipids and calculated long-range electrostatic interactions using the particle-mesh Ewald method (46). We set the switching length for electrostatics and the cut-off length for the Lennard-Jones potential at 10 Å. We treated water molecules and CHx, NHx, (x=1, 2, or 3), SH, and OH groups as rigid bodies using the LINCS (47) to allow a time step of 2.0 fs. We energy-minimized each simulation system using a 5000-step steepest descent method, followed by gradually decreasing restraints on protein and lipid heavy atoms for three consecutive 25 ps NVT simulations. In these simulations, we used a 1 fs time step and maintained a temperature of 303.15 K using a Berendsen temperature-coupling scheme. We then performed three consecutive 100 ps NPT simulations with further gradual release of heavy-atom restraints. In the last 100 ps simulation, the lipids were completely unrestrained and allowed to relax around the restrained protein. In these simulations, we used a 2 fs time step and kept the pressure constant at one bar using a semi-isotropic Berendsen pressure barostat. We performed independent 1 μs production runs once for FL and ΔSTAS and five times for ΔC.

## RESULTS AND DISCUSSION

### Role of STAS domain in ion transport

In this study, we performed seven independent 1-μs simulations of SLC26A9, including full-length (FL) × 1, STAS domain removal (ΔSTAS) × 1, and C-terminus removal (ΔC) × 5, in the presence of 150 mM NaCl under NPT ensemble conditions. We analyzed the probabilities of chloride ion present in the transmembrane region for each simulation and observed that in the FL (Figure 2a) and ΔSTAS (Figure 2b) simulations, no significant peaks in the probability of chloride ion presence were identified.

**FIGURE 2.**
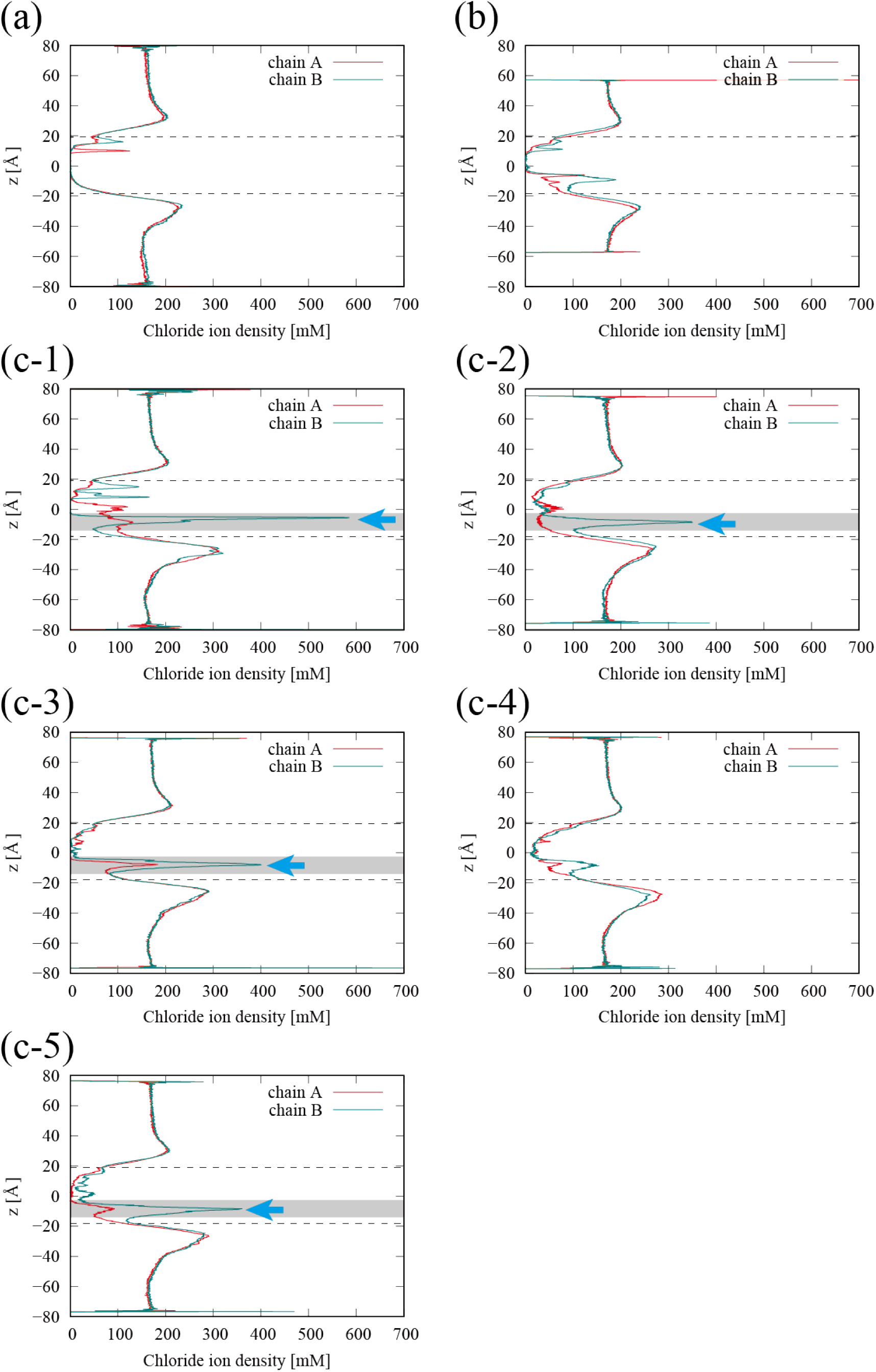
Distribution of the chloride ion density with respect to the direction perpendicular to the membrane (z). The red and green lines correspond to chloride ion densities in the chains A and B. The approximate locations of the phosphorus atoms in the headgroups of the lipid membranes are indicated by the dashed lines. The areas around the chloride ion binding sites are shaded, and the locations of the significant peaks in chloride ion densities are indicated by cyan arrows. (a) FL. (b) ΔSTAS. (c1-5) ΔC trajectory 1-5.

However, in ΔC simulations, a significant peak in the probability of the presence of chloride ions was observed in four out of the five simulations (Figure 2c 1-5). The chloride ion was found to be interacting with the main chains of THR127, PHE128, ALA390, LEU391, SER392, and side chains of GLN88, PHE128, ALA390, LEU391, SER392, ASN441, and ASN444, which are consistent with the cryo-EM structures of human SLC26A5 in the chloride binding state (27) (Figure 3).

**FIGURE 3.**
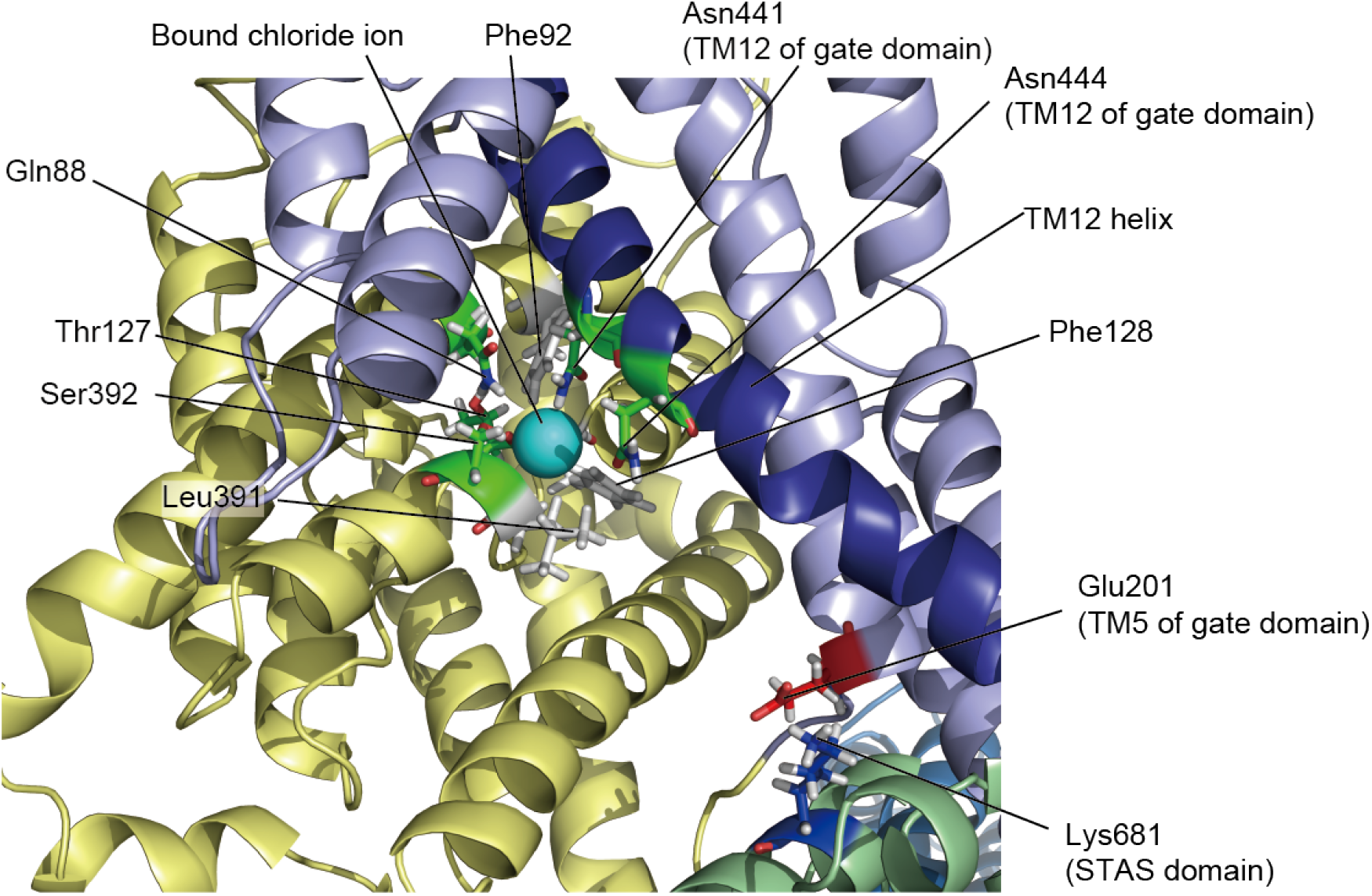
Snapshot of ΔC-t1 simulation at 750 ns. The colors of the cartoon represent the same elements as in Figure 1, with the exception of the TM12 helix of the gate domain which is colored deep blue. Stick representations of basic (blue), acidic (red), polar (green), and hydrophobic (white) residues entering in contact with the chloride ion or forming salt bridges that trigger the domain motion of STAS are shown. Chloride ion stably bound to the binding site is shown as a sphere (cyan).

The peak location of the probability of the presence of chloride ions is presumed to be the chloride ion binding site of SLC26A9. From the mutational analysis, it was revealed that the residues (GLN88, PHE92, THR127, PHE128, LEU391, SER392) surrounding the chloride ion have a significant impact on the transport characteristics of SLC26A9 (30). Importantly, the probability of the presence of chloride ions is almost zero at a position slightly closer to the extracellular side of the transmembrane domain than the binding site, indicating that the chloride ions entered the binding site from the cytoplasmic side and that these SLC26A9 structures are inward-facing open structures.

To transport chloride ions from the cytoplasmic side of the plasma membrane to the extracellular side in an elevator motion, chloride ions must be stably bound inside SLC26A9 for a long time while SLC26A9 changes from an inward open structure to an outward open structure. For trajectory 1 of ΔC, where the peak in the probability of chloride ion presence was observed to be most pronounced, the time variation of the probability of chloride ion binding to the binding site shown in Figure 4 and in Movie S1 indicates that stable ion binding has occurred over a long period of time. Therefore, the ΔC simulation demonstrated stable binding of chloride ions, which is necessary for transmembrane transport of ions, suggesting that the elevator motion mechanism operates.

**FIGURE 4.**
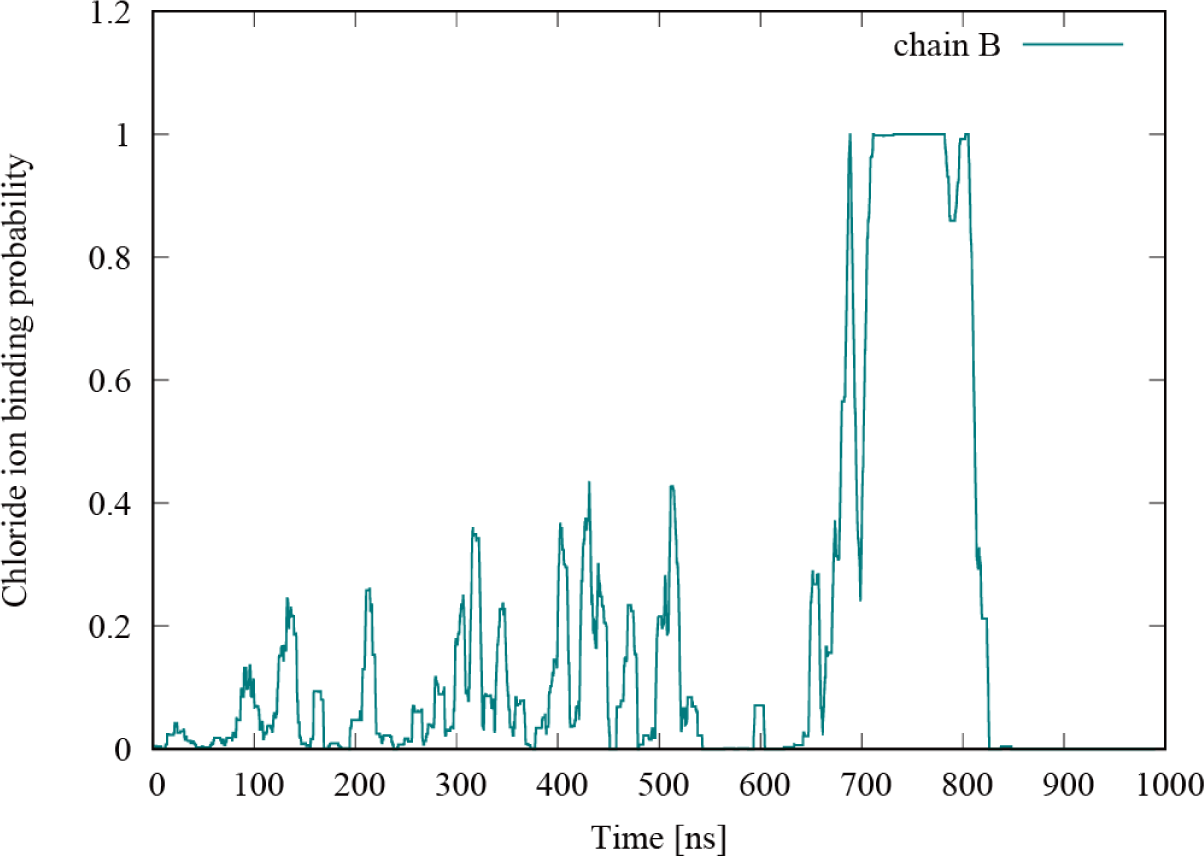
Time variation of the probability of chloride ion binding to the binding site of the chain B in ΔC-t1 simulation. Probabilities are window-averaged over a window width of 50 ns.

In summary, our simulations indicate that stable binding of chloride ions was observed only in ΔC, which may reflect the elevator motion mechanism. The C-terminal helix in the FL model plugged the ion pathway and inhibited chloride ions from accessing the binding site. Furthermore, the absence of stable binding of chloride ions in the ΔSTAS model suggests that the STAS domain is required for chloride ion transport. These observations are consistent with findings in previous studies (31,33-37). In the following studies, we further analyze the role of the STAS domain in regulating the access of chloride ions to the binding sites.

### Asymmetric motion of STAS domain promotes the gating of SLC26A9

Our simulations revealed that the transmembrane domains of the SLC26A9 dimer exhibit asymmetric behavior in the absence of the C-terminal helix, suggesting that the STAS domain plays a crucial role in maintaining the distance between the core domain and the gate domain. Figure 2 shows the chloride ions bound to binding sites in four of the five ΔC model simulations. However, in all those simulations, the chloride ion only accessed the binding site on one of the SLC26A9 dimers, indicating that the gating mechanism is asymmetric. Figure 5a-c presents the distribution of the distance between the core domain and the gate domain for the chains A and B in the FL, ΔSTAS, and ΔC-t1 simulations, respectively. The distance between the core domain and the gate domain was calculated as the distance between the Cα atoms of ASP362 and GLU201 located at the respective tips of the core and gate domains (Figure 5d). The distance in the initial structure was about 22 Å. In the FL simulation, the distances between the core domain and the gate domain did not change significantly from the initial structure for both the chains A and B (Figure 5a). On the other hand, in the ΔSTAS simulation, the distances between the core domain and the gate domain for both the chains A and B were smaller than in the initial structure (Figure 5b), indicating that the STAS domain helps to maintain the distance between the core and the gate domains.

**FIGURE 5.**
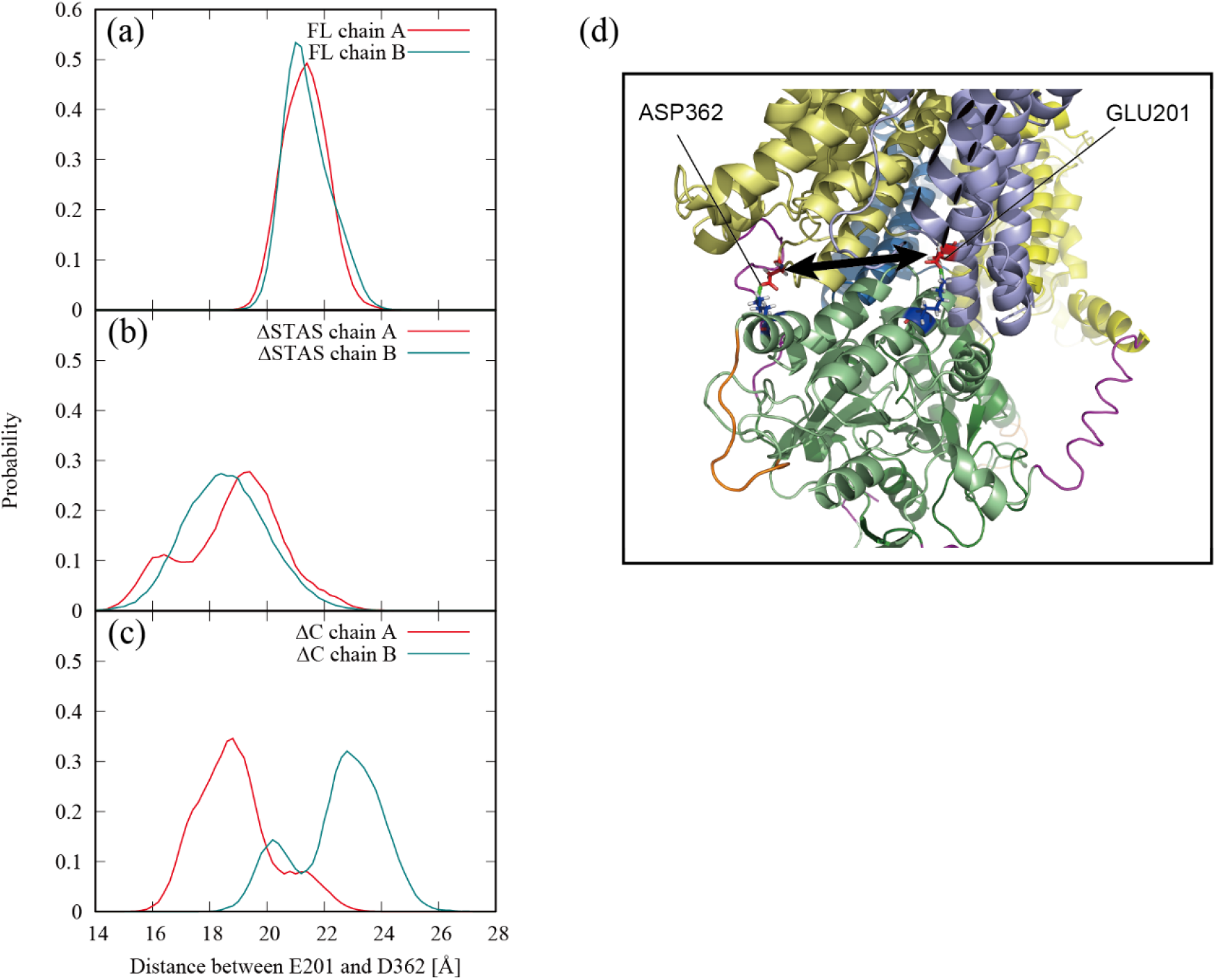
Distance between the core domain and the gate domain. Distribution of the distance between the core domain and the gate domain for the chains A (red) and B (green) in the (a) FL, (b) ΔSTAS, and (c) ΔC-t1 simulations. (d) Distance between the core domain and the gate domain was defined as the distance between the Cα atoms of ASP362 and GLU201. The colors of the cartoon represent the same elements as in Figure 1. Stick representation of basic (blue) and acidic (red) residues are also shown. Salt bridges are indicated by the green dotted lines.

Furthermore, the ΔC-t1 simulation showed asymmetric behavior, with the chain A having a smaller distance between the core and gate domains than the initial structure, while the chain B had a slightly larger distance than the initial structure (Figure 5c). This observation implies that not only the STAS domain but also the C-terminal helix affects the dynamic behavior of the transmembrane domain of SLC26A9. This difference in behavior may be attributed to the fact that the C-terminal helix plays a role in stabilizing the STAS domain and restricting its motion, as we will discuss next.

To better understand the role of the STAS domain in promoting the gating of SLC26A9, we analyzed the free energy landscape of the position of the center of mass of the STAS domain in the X-Y plane, viewed from the extracellular perspective, for the FL and ΔC-t1 simulations (Figure 6a-b).

**FIGURE 6.**
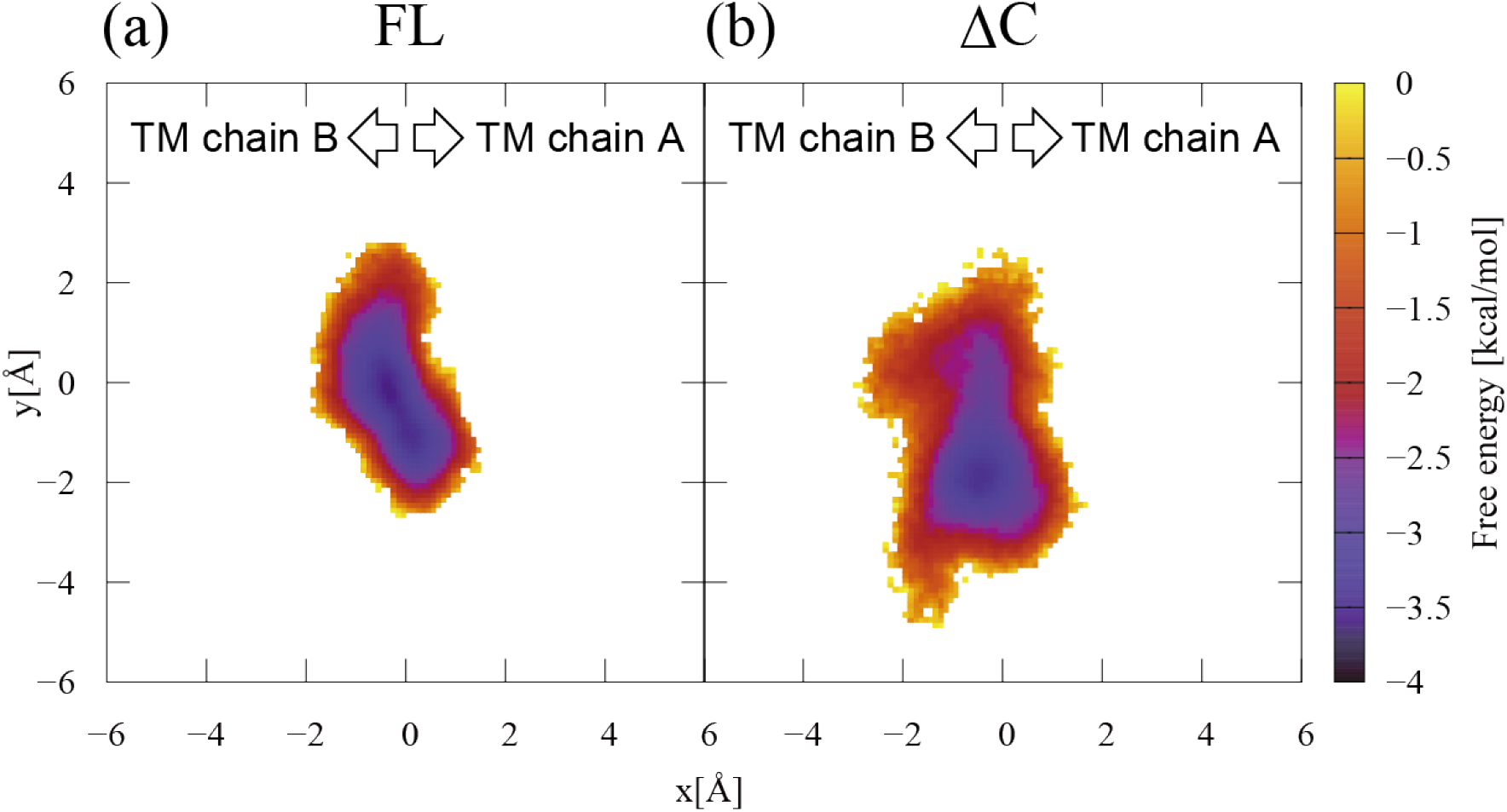
Free energy landscape of the position of the center of mass of the STAS domain for the (a) FL and (b) ΔC-t1 simulations. These landscapes are viewed from the extracellular perspective, with the transmembrane domains of the chains A and B located on the right and left sides, respectively.

We found that the STAS domain of ΔC has a larger and more asymmetric motion than the STAS domain of FL. In FL, the PDZ-binding motif in the C-terminal region is stuck between the GLU201 in the gate domain and LYS681 in the STAS domain (Figure 7a), thus restricting the movement of the STAS domain. In contrast, in the case of ΔC, the deletion of the C-terminal helix is thought to create a salt bridge between GLU201 and LYS681, which may excite the motion of the STAS domain. This large and asymmetric motion of the STAS domain of ΔC resulted in different salt bridge formations between the chains, as we discuss next.

**FIGURE 7.**
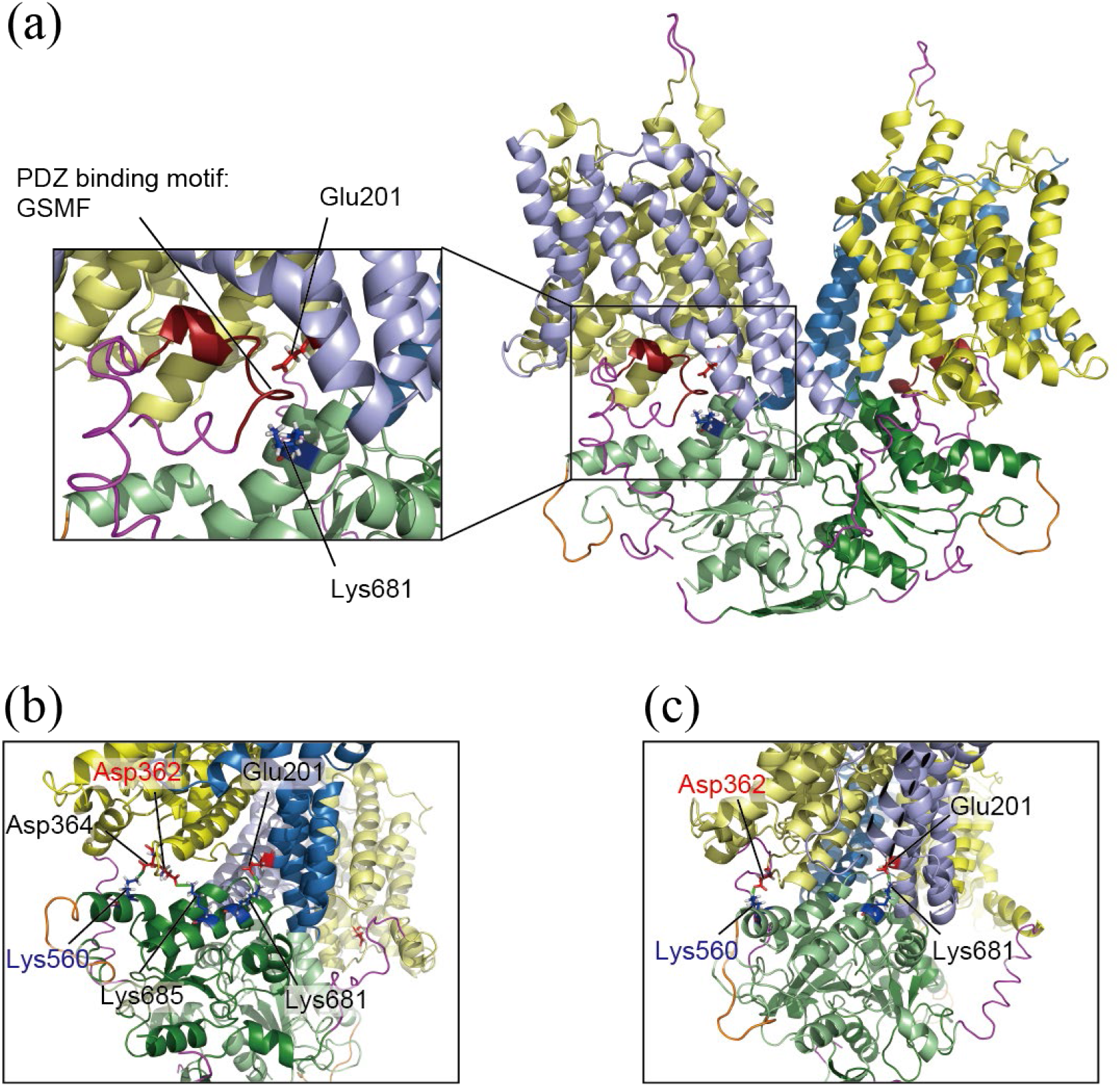
Different salt bridge formations between the chain A and the chain B resulted by the deletion of the C-terminus. (a) Close up view around the C-terminal helix of the SLC26A FL model. (b) Snapshot from (b) the chain A side and (c) the chain B side of ΔC-t1 simulation at 664 ns. The colors of the cartoon represent the same elements as in Figure 1. Stick representation of basic (blue) and acidic (red) residues are also shown. Salt bridges are indicated by the green dotted lines.

In the ΔC-t1 simulation, in the transmembrane domain of chain A, ASP362 in the core domain of chain A formed a salt bridge with LYS685 in the STAS domain of chain B and ASP364 in the core domain of chain A formed a salt bridge with LYS560 in the STAS domain of chain B (Figure 7b), whereas in the transmembrane domain of chain B, ASP362 in the core domain of chain B and LYS560 in the STAS domain of chain A formed a salt bridge (Figure 7c). This difference in salt bridge formation may have resulted in the difference in the relative positions of the core and gate domains between the chains A and B. Figure 8 shows the free energy landscape of chloride ion binding versus the distance between the core and gate domains, calculated from five ΔC simulations. This free energy landscape means that the binding of chloride ions favors a structure in which the distance between the core and gate domains is slightly larger than in the initial structure. The increase in the distance between the core domain and the gate domain corresponds to the gating of the pathway through which chloride ions are transported.

**FIGURE 8.**
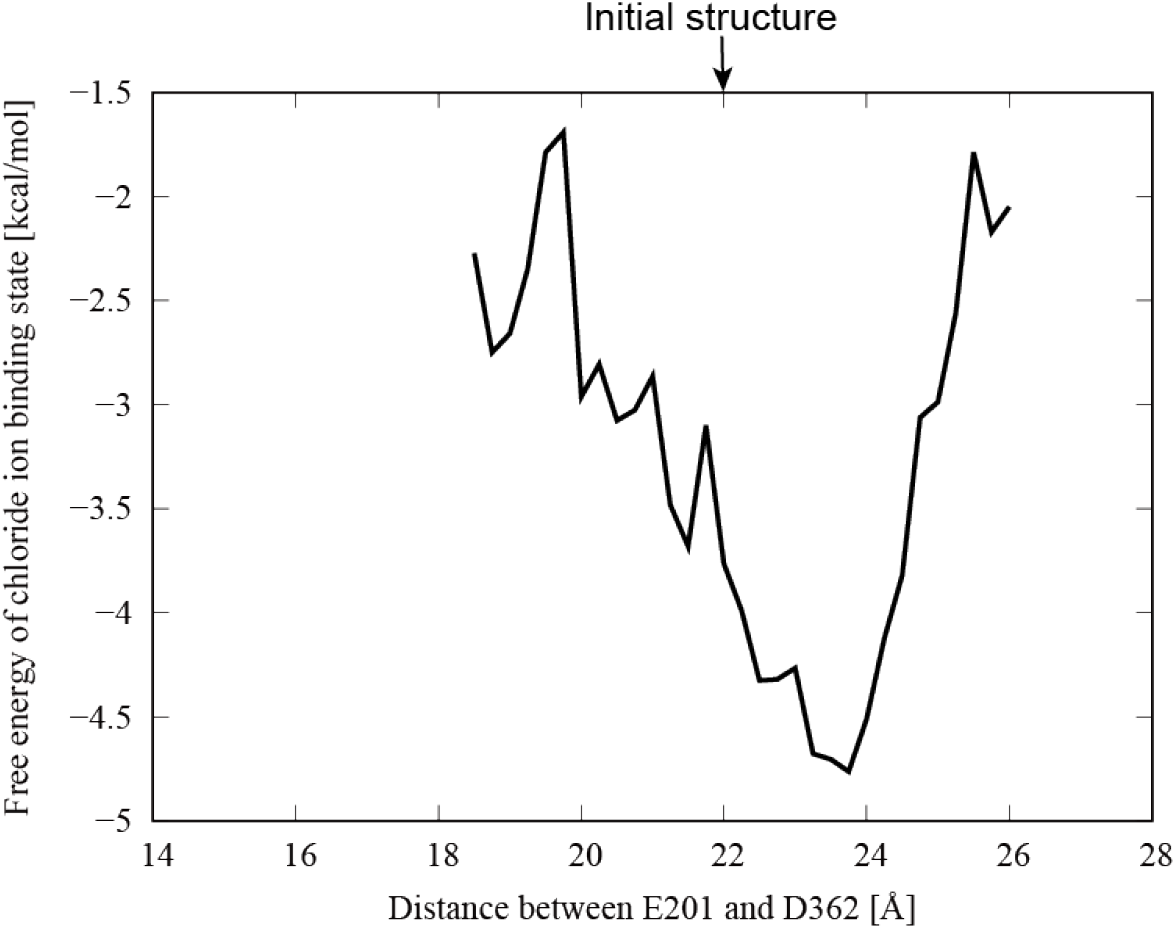
Free energy landscape of chloride ion binding versus the distance between the core and gate domains calculated from five ΔC simulations.

To summarize, our findings suggest that the removal of the C-terminal helix triggers asymmetric motion of the STAS domain, which ultimately opens the gate of the transmembrane domain and enables chloride ions to bind to the binding site. However, it remains unclear how ΔC stably binds chloride ions to the binding site. In the following section, we analyze how the STAS domain participates in the elevator motion of the transporter.

### Transformation of the TM12 helix stabilizes the binding of chloride ion

In this section, we discuss how the transformation of the TM12 helix stabilizes the binding of chloride ion in the ΔC simulation. The TM12 helix is a gate domain helix that interacts with the C-terminal helix to form a kinked structure in the cryo-EM structure. In the initial structure, LYS447 and GLU775 interacted by a salt bridge (Figure 9a). In the ΔC simulation, the asymmetric motion of the STAS domain leads to the formation of a salt bridge between the TM12 and STAS domains (Figure 9b). This salt bridge formation is also observed in the time variation of the distance between the Nζ atom of LYS458 at the end of the TM12 helix of the chain B and the Oδ atom of ASP709 at the interface of the STAS domain of the chain A in the ΔC-t1 simulation (Figure 9c).

**FIGURE 9.**
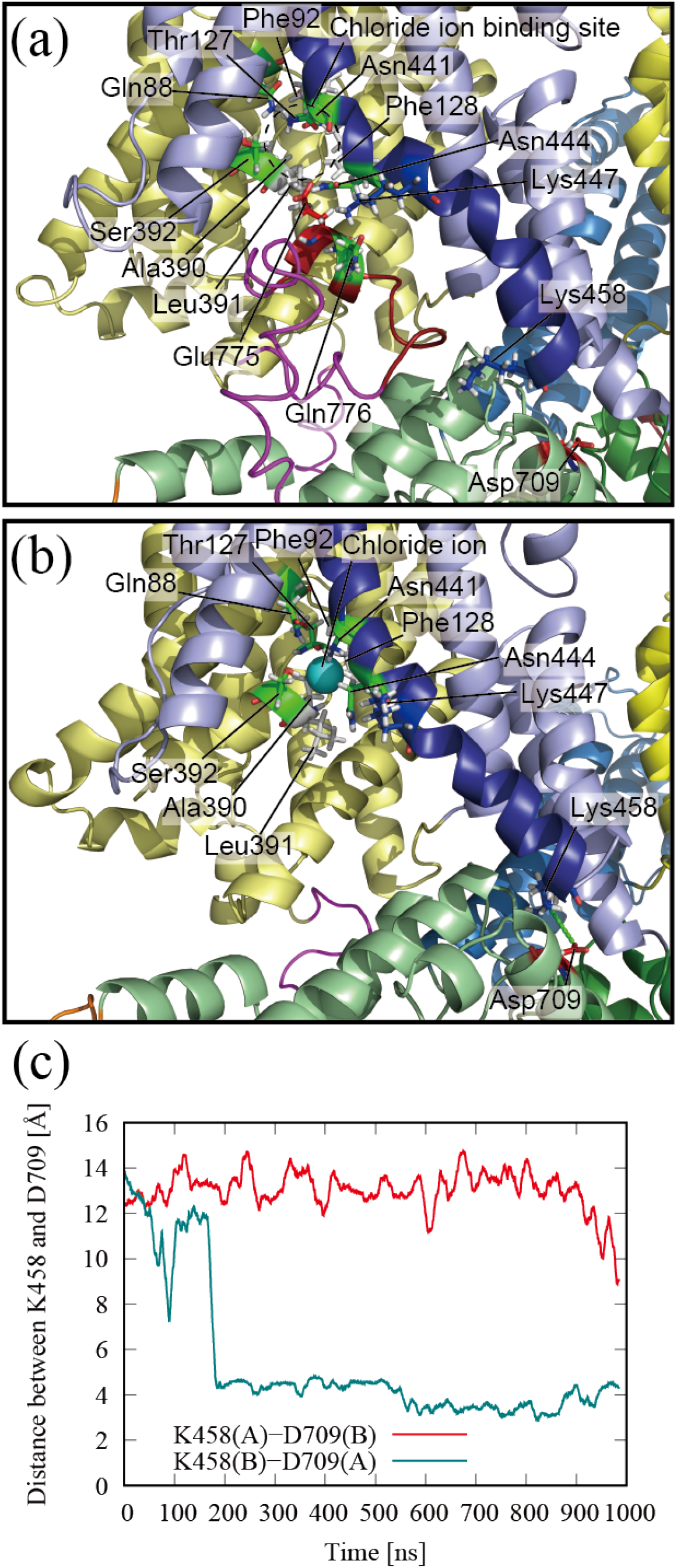
Transformation of the TM12 helix by the deletion of the C-terminus. (a) Close up view around the C-terminal helix of the SLC26A9 cryo-EM structure (PDB ID: 7CH1) and (b) snapshot of the same view at 750 ns. The colors of the cartoons and sticks in panels (a) and (b) have the same signification as in Figure 3, with the C-terminal ordered and disordered regions shown in dark red and magenta, respectively. Stick representation of basic (blue), acidic (red), polar (green), and hydrophobic (white) residues are also shown. (c) Time variation of distance between the Nζ atom of LYS458 at the end of the TM12 helix of the chain B and the Oδ atom of ASP709 at the interface of the STAS domain of the chain A in the ΔC-t1 simulation.

The clearance between the core domain and the gate domain, created by the removal of the C-terminal helix and the large motion of the STAS domain, enables the transformation of the TM12 helix from its initial kinked structure. Figure 10a quantifies the transformation of the TM12 helix in the ΔC-t1 simulation. The helix kink is calculated as the angle between the helix axis segments. ASN444 and SER445 in the TM12 helix of the chain B were not kinked in the initial structure, but kinked significantly at 200 ns, coinciding with the formation of a salt bridge between LYS458 (B) and GLU709 (A) (Figure 10a top). At the same time, the main chain dihedral angle ASN444ψ (B) also changed significantly (Figure 10a middle). However, at this point in chain B, there is no significant change in the distance between LEU128, which is in the core domain and forms a chloride ion binding site, and ASN444 of the TM12 helix (Figure 10a bottom). Therefore, the transformation of the TM12 helix at 200 ns, due to the change in kink and dihedral angles, is not considered to be involved in stabilizing chloride ion binding. Subsequently, under the influence of STAS motion, the kink angles of ASN444 and SER445 in the chain B become smaller again at 500 ns (Figure 10a top), and at the same time, ASN444ψ changes significantly (Figure 10a middle). From this 500 ns point, the distance between LEU391 and ASN444 in chain B becomes smaller (Figure 10a bottom).

**FIGURE 10.**
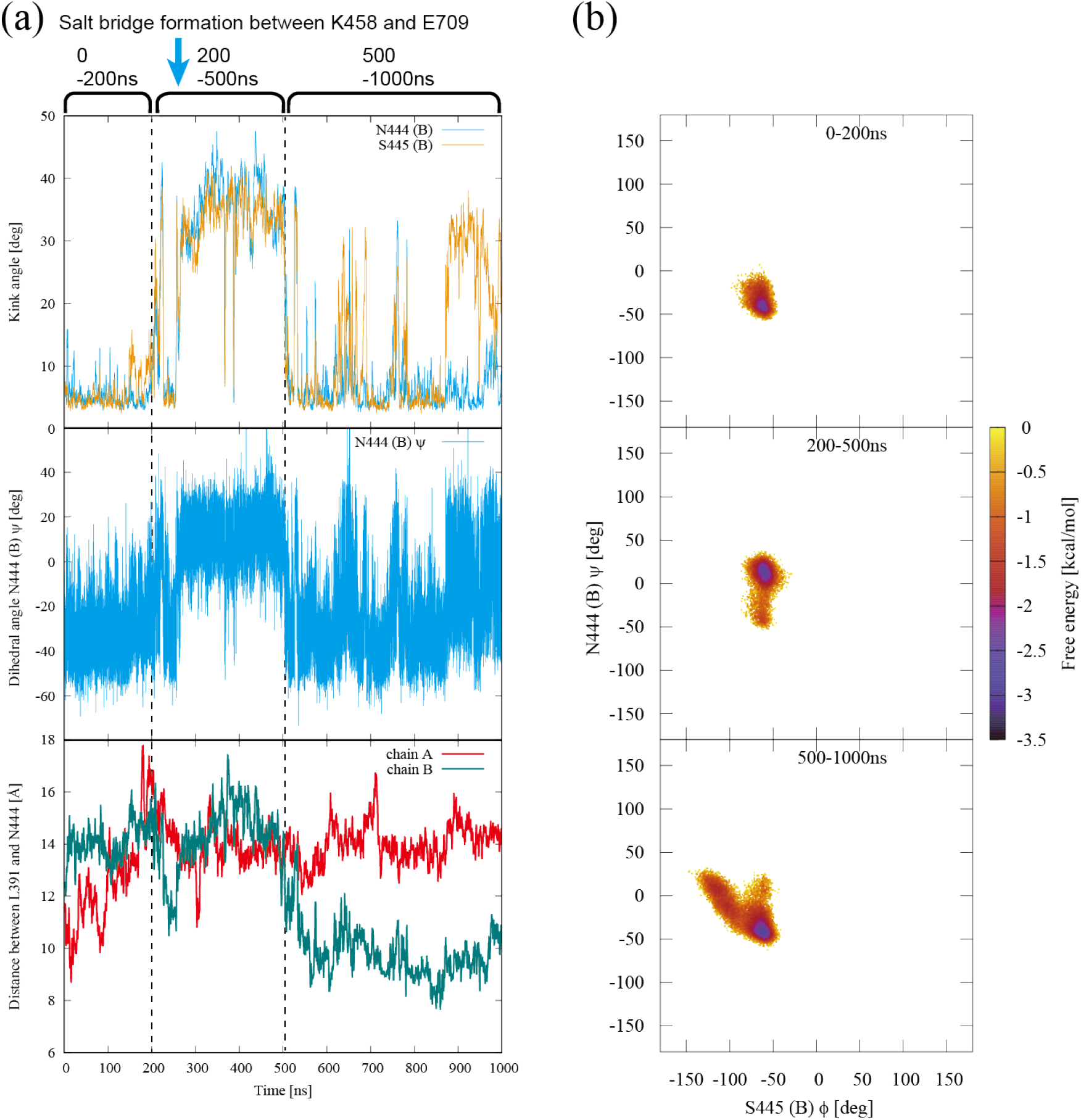
Stiffening of the TM12 helix to stabilize the ion binding. (a) Quantitative values for the deformation of the TM12 helix in the ΔC-t1 simulation. The time point when the salt bridge is formed between LYS458 (chain B) and GLU709 (chain A) is indicated by the cyan arrow. The time point when large changes in kink angles occur is indicated by the dashed line. (Top) Time variation of kink angle for ASN444 (chain B) (cyan) and SER445 (chain B) (orange). (Middle) Time variation of main chain dihedral angle ASN444ψ (chain B). (Bottom) Time variation of the distance between the Cα atoms of LEU391 and ASN444 in chains A (red) and B (green). (b) Free energy landscape showing the correlation between the dihedral angles SER445φ and ASN444ψ of the chain B in the ΔC-t1 simulation. (Top) 0- 200 ns. (Middle) 200-500 ns. (Bottom) 500-1000 ns.

The smaller distance between LEU391 and ASN444 facilitates the formation of clusters of hydrophilic amino acids around the chloride ion binding sites of the core domain, as shown in Figure 3. This configuration allows for the stable binding of chloride ions, as depicted in Figure 4. Therefore, the transformation of the TM12 helix at 500 ns, accompanied by changes in kink angle and dihedral angle, played a significant role in stabilizing the binding of chloride ions. These results indicate that the TM12 helix exhibited distinct properties at 200 ns and 500 ns. Figure 10b presents a free energy landscape that reveals the correlation between the dihedral angles SER445φ and ASN444ψ in the ΔC-t1 simulation. The changes in both SER445φ and ASN444ψ during the 0-200 ns interval were small, suggesting that the structure of the TM12 helix remained relatively unchanged (Figure 10b top). However, after ASN444 and SER445 significantly kinked due to the formation of the salt bridge between LYS458 (chain B) and GLU709 (chain A) at 200 ns (Figure 10a top), ASN444ψ changed significantly in an uncorrelated manner with SER445φ during 200-500 ns (Figure 10b middle). The uncorrelated change in dihedral angle implies that the structure of the TM12 helix was flexible during this period, and therefore, the cluster of hydrophilic amino acids surrounding the chloride ion binding site may not have formed. Conversely, after the kink angle between ASN444 and SER445 became small again at 500 ns (Figure 10a top), ASN444ψ and SER445φ changed in an inversely correlated manner (Figure 10b bottom). This motion is known as crankshaft motion (48), which maintains the overall structure of the protein despite the large dihedral displacement. In other words, crankshaft motion imparts stiffness to the structure. The TM12 helix exhibits a stiff structure, which is thought to be pulled by the movement of the STAS domain toward the chloride ion binding site (Figure 10a bottom). This, in turn, promotes the formation of the cluster of hydrophilic amino acids surrounding the chloride ion binding site, stabilizing the binding of chloride ions (Figure 3 and Figure 10a bottom).

The dihedral angle motion of the TM12 helix in SLC26A9 depends on the angle between the main chains of adjacent residues. In other words, the TM12 helix switches the dihedral correlation between non-crankshaft and crankshaft modes by changing the shape of the kink. Table 1 shows the relationship between the dihedral angular correlation of TM12 and the presence or absence of stable binding of chloride ions for each chain in the simulations in this study. The correlation of the dihedral angles with both non-crankshaft and crankshaft correlations is denoted as dual mode, while the correlation with only non-crankshaft correlations is denoted as single mode.

**TABLE 1.**
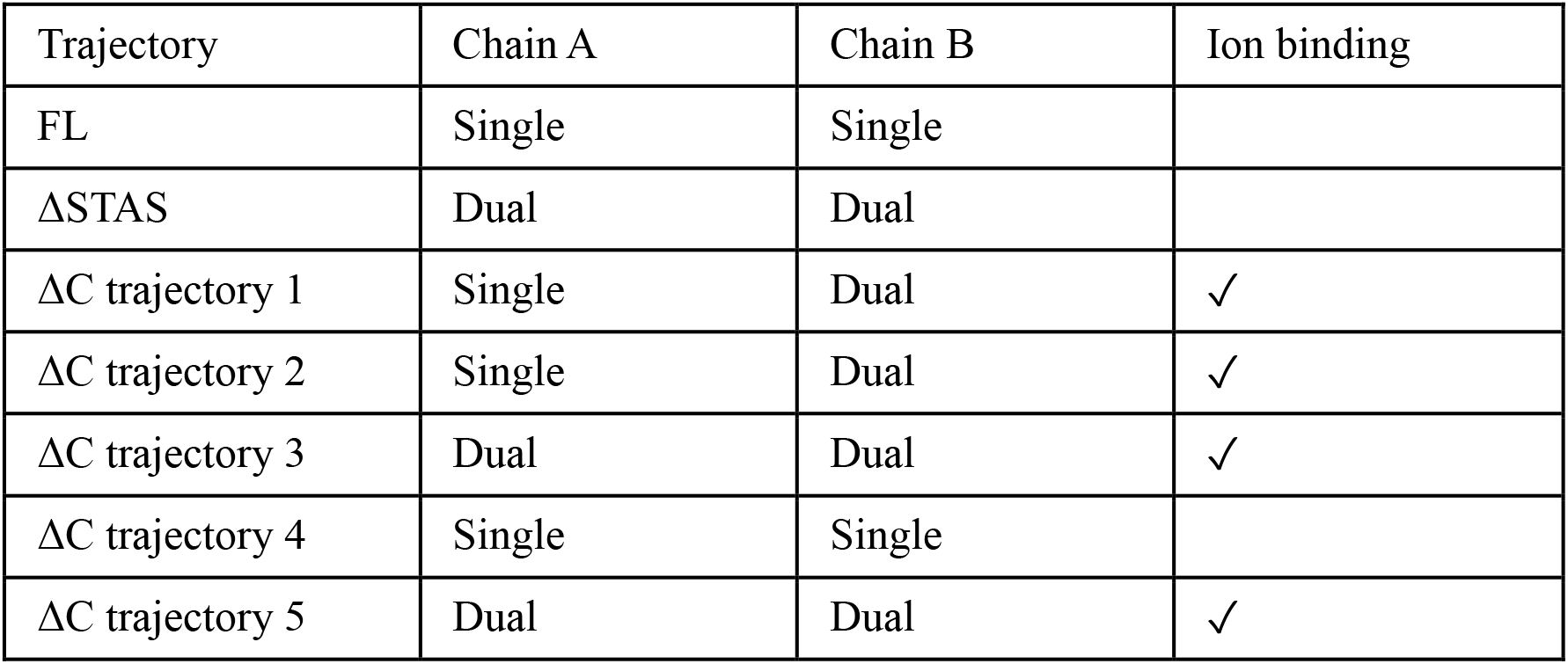
The relationship between the dihedral angular correlation of TM12 and the stable binding of chloride ions.

| Trajectory | Chain A | Chain B | Ion binding |
| --- | --- | --- | --- |
| FL | Single | Single |  |
| ΔSTAS | Dual | Dual | |
| ΔC trajectory 1 | Single | Dual | ✓ |
| ΔC trajectory 2 | Single | Dual | ✓ |
| ΔC trajectory 3 | Dual | Dual | ✓ |
| ΔC trajectory 4 | Single | Single | |
| ΔC trajectory 5 | Dual | Dual | ✓ |

The TM12 helix in the chains where chloride ion binding was observed had dual mode dihedral correlations. This suggests that the dual mode dihedral angular correlation brought about by the kinked structure of the TM12 helix is required for stable binding of chloride ions.

Combining the results so far in this study and knowledge from existing studies, the elevator motion of SLC26A9 can be interpreted as shown in the schematic view in Figure 11. Before insertion of SLC26A9 into the apical plasma membrane, the C-terminal helix is in the chloride ion pathway between the core and gate domains (Figure 11a). Although the sequence of the C-terminal region in the SLC26 family is not well conserved, some SLC26 transporters harbor a PDZ domain binding motif at their C-termini. SLC26A9 has an X–S/T–X–Φ of class I type PDZ domain binding motif, where Φ represents a hydrophobic residue, and interacts with the PDZ domain of the scaffold protein NHERF1. From existing research, after insertion of SLC26A9 into the apical plasma membrane, its C-terminus is expected to dissociate from the intracellular vestibule and interact with NHERF1 (Figure 11b). The salt bridge between the gate domain and the STAS domain is created by the dissociation of the C-terminal helix, thereby triggering the asymmetric motion of the STAS domain (Figure 11c). Asymmetric motion of the STAS domain causes gating that increases the distance between the core domain and the gate domain, allowing chloride ions to enter their binding site (Figure 11d). In addition, the TM12 helix interacts with the STAS domain by the salt bridge and transforms the kink structure into a stiff structure and forms hydrophilic clusters for stable binding of chloride ions (Figure 11e). If SLC26A9 changes to an outward open conformation while chloride ions remain bound to the binding site, it would be able to transport chloride ions from the cytoplasm to the extracellular environment.

**FIGURE 11.**
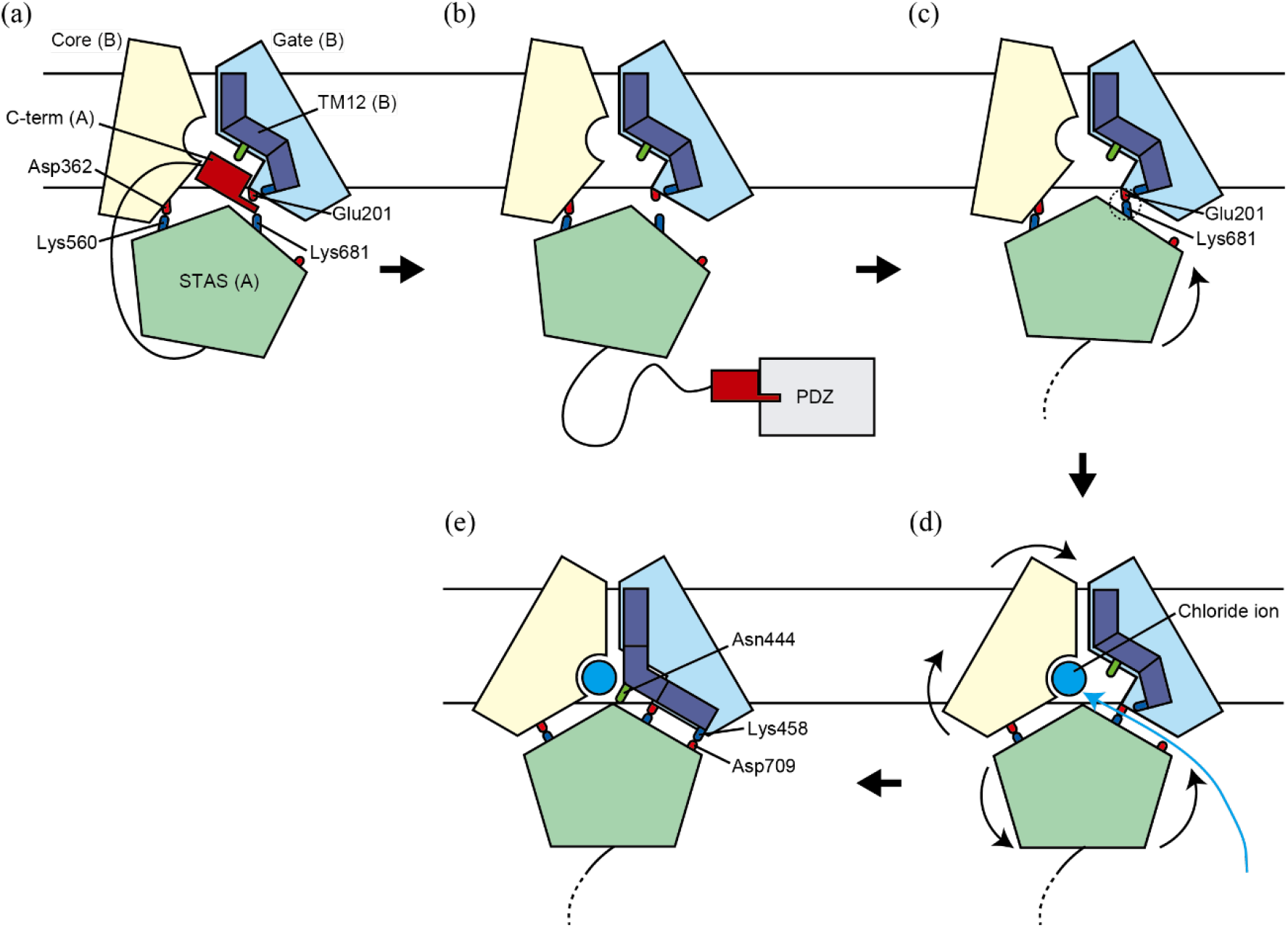
Schematic view of elevator motion of SLC26A9. The viewing angle is based on that on the left in Figure 1a. Core domain, gate domain, and TM12 helix of chain B are shown in pale yellow, light blue, and deep blue, respectively. The C-terminal helix and the STAS domain of chain A, the PDZ domain of the scaffold protein, and the chloride ion are shown in dark red, light green, gray, and cyan, respectively. Basic, acidic, and polar residues are shown as blue, red, and green sticks, respectively. Salt bridges are represented by dashed circles. Locations of the headgroups of the lipid membranes are indicated as the black solid lines. (a) Before insertion of SLC26A9 into the apical plasma membrane. (b) C-terminus dissociates from the intracellular vestibule and interacts with the PDZ domain of the scaffold protein after insertion of SLC26A9 into the apical plasma membrane. (c) Salt bridge between the gate domain and the STAS domain is created by the removal of the C-terminal helix. (d) Distance expansion between core and gate domains caused by asymmetric motion of the STAS domain, allowing chloride ions to enter their binding site. (e) Transformation of TM12 by contact with STAS domain forms hydrophilic clusters for stable binding of chloride ions.

## CONCLUSION

This study sheds light on the elevator motion mechanism in the chloride ion transporter SLC26A9. We found that removing the C-terminal helix not only opens the ion pathway but also initiates STAS domain motion, leading to asymmetric gating of the transmembrane domain and stiffening of the flexible helix near the ion binding site. This structural change allows stable chloride ion binding, essential for elevator motion.

Our findings lay a robust groundwork for further research into the ion transport mechanisms of SLC family proteins. An enhanced understanding of SLC26A9’s elevator motion could influence the development of new therapeutics targeting these proteins.

In conclusion, we have clarified how the STAT domain and TM12 helix contribute to stabilizing chloride ion binding in the SLC26A9 transporter. The transformation of the TM12 helix and its interaction with the STAS domain form hydrophilic clusters that facilitate stable ion binding. These insights not only enrich our comprehension of SLC26A9’s function but also offer potential guidance for therapeutic strategies addressing diseases related to chloride ion transport dysfunction.

## AUTHOR CONTRIBUTIONS

The contributions of each author to this work are as follows: HN conceptualized and proposed the idea for the research study. YH was responsible for selecting the subjects. SO performed all calculations and was heavily involved in the data analysis, along with all the other authors. SO and YH also took the lead in writing the manuscript. KK provided overall supervision, revising the manuscript, ensuring coherence and integrity in the study, securing the budget, and managing the project.

## ACKNOWLEDGEMENTS

This research was supported by Canon Medical Systems Corporation and the Platform Project for Supporting Drug Discovery and Life Science Research (Basis for Supporting Innovative Drug Discovery and Life Science Research (BINDS)) from Japan Agency for Medical Research and Development (AMED) under Grant Number JP21am0101067 and by the Research Support Project for Life Science and Drug Discovery (BINDS) from AMED under Grant Number JP22ama121019. All computational resources were provided by the ToMMo supercomputer system (http://sc.megabank.tohoku.ac.jp/en).

